# Comparative genomics of Minnesotan barley-infecting *Xanthomonas translucens* shows overall genomic similarity but virulence factor diversity

**DOI:** 10.1101/2022.04.05.487151

**Authors:** Nathaniel Heiden, Verónica Roman-Reyna, Rebecca Curland, Ruth Dill-Macky, Jonathan M. Jacobs

## Abstract

*Xanthomonas translucens* pv. translucens (Xtt) is a global barley pathogen and a concern for resistance breeding and regulation. Long-read whole genome sequences allow in-depth understanding of pathogen diversity. We have completed long-read PacBio sequencing of two Minnesotan Xtt strains and an in-depth analysis of available Xtt genomes. We found that average nucleotide identity(ANI)-based approaches organize Xtt strains differently than the previously standard MLSA approach. According to ANI, Xtt forms a separate clade from *Xanthomonas translucens* pv. undulosa and consists of three main groups which are represented on multiple continents. The global distribution of Xtt groups suggests that regulation of seed is not important for prevention of Xtt spread. Some virulence factors, such as 17 Type III-secreted effectors, are highly conserved and offer potential targets for the elicitation of broad resistance. However, there is a high degree of variation in virulence factors meaning that germplasm should be screened for resistance with a diverse panel of Xtt.

---

*Xanthomonas translucens* pathovar (pv.) translucens (Xtt) causes bacterial leaf streak, blight and black chaff of barley (Jones et al. 1917; Bragard et al. 1997; Sapkota et al. 2020). Xtt foliar infections of barley cause translucent streaks that develop into necrotic lesions. Xtt exudes out of watersoaked lesions providing an opportunity for short distance dispersal. Foliar infections are called bacterial leaf streak due to their symptomology and in severe cases are referred to as bacterial blight. Black chaff is a disease of the grain heads characterized by darkening of the glumes that is associated with seed transmission of Xtt. Little is known about the seed infection process, how black chaff develops or if seed infection is the main mechanism of Xtt dispersal. Xtt is increasingly impactful to cereal growers worldwide with recent widespread reports in the northern United States (Curland et al. 2020), Iran (Habibian et al. 2021) and Canada (Tambong et al. 2021). Though losses in barley due specifically to Xtt have not been quantified, wheat farmers can experience yield reduction as high as 40% due to infection with the closely related pathogen *X. translucens* pv. undulosa (Xtu) (Forster and Schaad 1988). There is no available single gene resistance to Xtt in barley breeding programs or for commercial growers.

Xtt is classified in the genomic subgroup Xt-I (Sapkota et al. 2020; Goettelmann et al. 2022), which also includes wheat and barley-infecting Xtu. Xtt was historically divided into three groups (A, B and C) according to multilocus sequencing analysis (MLSA) of four housekeeping genes (Curland et al. 2018). Strains from these three groups are present globally (Curland et al. 2018; Roman-Reyna et al. 2020; Shah et al. 2021). It is unknown if average nucleotide identity (ANI) based on whole genome analyses would confirm the phylogenetic groups proposed by MLSA. Recent research has also provided insights into virulence factor diversity through draft and whole genome analysis of some Xtt isolates (Peng et al. 2016; Langlois et al. 2017; Roman-Reyna et al. 2020; Shah et al. 2021; Jaenicke et al. 2016). These analyses have enabled the development of diagnostic primers for general *X. translucens* identification (Langlois et al. 2017) and also strengthened our understanding of virulence factor repertoires from Asian Xtt (Roman-Reyna et al. 2020; Shah et al. 2021).

The only strain from the Western Hemisphere with a publicly available long-read genome was isolated in 1933 (Jaenicke et al. 2016). The lack of genome resources is a roadblock for defining North American Xtt virulence factors or immune elicitors for barley resistance screening. A better definition of North American Xtt genomic composition and virulence factor prevalence will directly inform breeders on the most desirable host factors for conferring durable resistance to Xtt. This is also important background information for identification efforts to understand pathogen dispersal and survival and may have implications for regulations that have been based on isolates from outside of North America.

In this study we generated and analyzed high quality, complete genomes of two Xtt strains: CIX43 and CIX95. We previously characterized both strains as pathogenic on barley and non-pathogenic on wheat (Curland et al. 2018). CIX43 and CIX95 were isolated from barley in Minnesota in 2009 and 2011 and are representative of Xtt MLSA groups A and C, respectively. These strains were previously included in diversity analyses and serve as a reference for strains currently used in resistance screening programs. Therefore, we characterized these genomes to enhance diversity analysis and to help define how virulence factors relate to the broad Xtt population diversity.

DNA was extracted with the Genomic DNA Buffer Set and Genomic-tip 100/G (QIAGEN®) and sequenced, in 2019, with PacBio RSII (P6-C4) and 20kb SMRT bell library (Psomagen, Rockville, MD). Reads were assembled with Flye version 2.4 (Kolmogorov et al. 2019) and genome assemblies were annotated with the National Center for Biotechnology Information (NCBI) Prokaryotic Genome Annotation Pipeline (Tatusova et al. 2016) and are publicly available (Table 1). GC content was calculated using the Rapid Annotation of microbial genomes using Subsystems Technology (Overbeek et al. 2014).

**Table 1.** *Xanthomonas translucens* genomes* analyzed in this study.

| Strain – Accession | Publication | Year Isolated | Country | Genome size | Contigs | ANI | Secreted Proteins |
| --- | --- | --- | --- | --- | --- | --- | --- |
| CIX43 – CP072988;CP072989 | This study | 2009 | USA | 4.70 | 2 | 99.55 | 752 |
| CIX95 – CP072990 | This study | 2011 | USA | 4.65 | 1 | 99.26 | 731 |
| UPB886 – GCA_009600865.1 | Roman-Reyna et al. 2020 | 1990 | Iran | 4.67 | 2 | 98.94 | 702 |
| DSM 18974 - LT604072 | Jaenicke et al. 2016 | 1933 | USA | 4.72 | 1 | 100.00 | 761 |
| XtKm7 - CP064005 | Shah et al. 2021 | 2014 | Iran | 4.58 | 1 | 99.52 | 710 |
| XtKm8 - CP064004 | Shah et al. 2021 | 2014 | Iran | 4.79 | 1 | 99.10 | 751 |
| XtKm9 - CP064003 | Shah et al. 2021 | 2015 | Iran | 4.69 | 1 | 98.92 | 709 |
| XtKm34 - CP064001 | Shah et al. 2021 | 2015 | Iran | 4.68 | 1 | 99.11 | 731 |
| B2 - GCA_001542205.1 | Peng et al. 2016 | 2013 | USA | 4.54 | 517 | 99.00 |  |
| UPB787 - GCA_001469515.1 | Bragard et al. 1997 | 1990 | Paraguay | 4.54 | 763 | 98.97 |  |
| XT8 - GCA_001462125.1 | Peng et al. 2016 | 1942 | Canada | 5.71 | 156 | 98.98 |  |
| UPB458 - GCA_001659915.1 | Bragard et al. 1997 | 1970 | India | 4.52 | 1,156 | 99.00 |  |
| B1 - LNTA000000000.1 | Peng et al. 2016 | 2013 | USA | 4.80 | 523 | 99.09 |  |
| LW16 - GCA_001462075.1 | Peng et al. 2016 | 2009 | USA | 4.69 | 475 | 97.61 |  |
| CFBP 2541 - CP074364 | Pesce et al. 2015 | 1941 | USA | 4.50 | 1 | 94.94 |  |

\*Genome summary and predicted secreted proteins of *X. translucens* pathovar translucens. Genome size and CDS data were determined with PGAP analysis (Tatusova et al. 2016) for strains CIX43 and CIX95 and publicly available information from NCBI was used for other strains. ANI is relative to type *X. translucens* pv. translucens strain DSM 18974, determined by Enveomics (Rodriguez-R and Konstantinidis 2016).

The CIX43 genome is 4,700,914 base pairs and has a total of 3,990 coding sequences (CDS). It contains two circular contigs of 4,664,501 bp and 36,413 bp with mean coverages of 141X and 53X, respectively. This translates to an N_50_ of 4,664,501 and L_50_ of 1. The second contig is a plasmid based on the results from Blastn search against the NCBI nucleotide collection database (Zhang et al. 2000). The results indicate the plasmid has 81% coverage and 85.92% identity to a X. campestris pv. campestris plasmid. No significant homology to *X. translucens* genomes were found in the same search. The CIX95 genome has 4,647,206 base pairs in a single contig with a mean coverage of 173X and 3,926 total CDS, for an N_50_ of 4,647,206 and L_50_ of 1. Both strains have a GC content of 67.8% and their average nucleotide identity (ANI) is 99.24% (Table S1).

The geographic distribution of Xtt populations remains unclear. Xtt has been isolated from all continents except Antarctica (Sapkota et al. 2020), but it remains uncertain if this distribution is from seed or an unknown environmental source. Phylogenomics provides a method to capture Xtt genetic diversity and characterize Xtt subgroups to begin to infer inoculum sources. To define the genomic relationships among CIX43, CIX95 and 11 additional Xtt genomes (Table 1), ANI and life identification numbers (LINs) were calculated with the webtools Enveomics and LINbase, respectively (Rodriguez-R and Konstantinidis 2016; Tian et al. 2020). Xtu LW16 and *X. translucens* pv. cerealis (Xtc) CFBP 2541 were used as outgroups (Pesce et al. 2015).

The Enveomics ANI and LINbase LIN analyses demonstrate that there are three major Xtt groups, which are internationally dispersed and distinct from Xtu and Xtc (Fig. 1). CIX43 and CIX95 are in the same phylogenetic cluster and LINgroup. Xtt strains have a high degree of homology as they share at minimum 98.88% ANI (Table S1). Analyzed Xtt strains have between 97.56% and 97.79% ANI to Xtu LW16 and between 94.89% and 95.07% ANI to Xtc strain CFBP 2541 (Table S1). We further validated this approach by providing LINs for each of the analyzed strains. According to their LINs, Xtt strains form a separate LINgroup (15_A_,1_B_,1_C_,0_D_,0_E_,0_F_,1_G_,0_H_,0_I_,0_J_) from Xtu and Xtc are best divided into three LIN subgroups with a minimum of 99% ANI within each: 1(Xtt:1_K_), 2(Xtt:0_K_), 3(Xtt:2_K_) (Fig. 1). Strains from different years and locations intermixed, which argues against seed dissemination by these pathogens.

**Figure 1.**
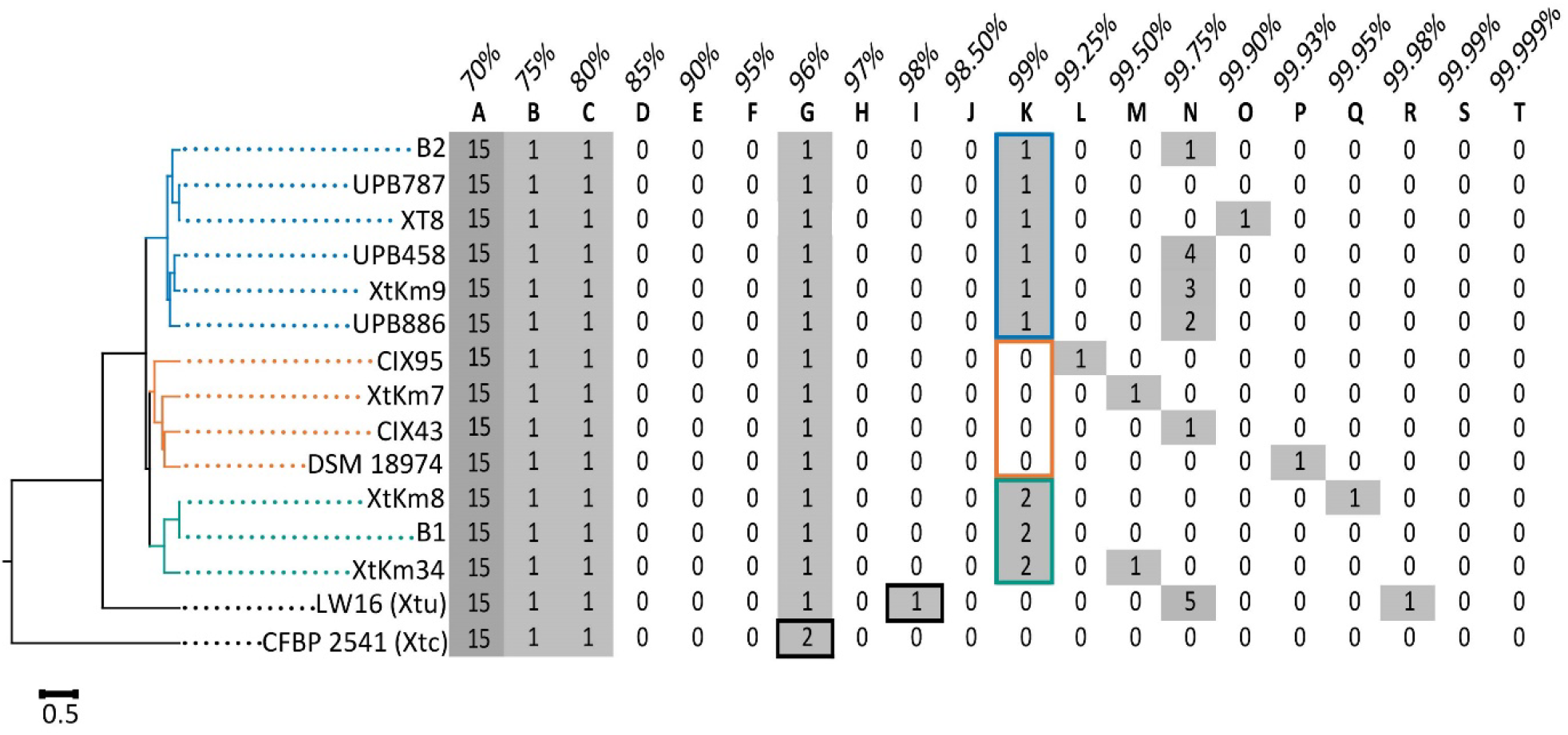
Xtt strains are separate from Xtu and form three distinct phylogenetic groups according to ANI. Whole genome ANI was calculated for all publicly available *X. translucens pv. translucens* strains and the outgroup strains LW16 and CFBP 2541. A tree was generated with the webtool Enveomics using the UPGMA clustering method (Rodriguez-R and Konstantinidis 2016). Life identification numbers were calculated with LINbase (Tian et al. 2020). Boxes outline the LIN values which separate the subgroups.

Traditional diversity analysis for *Xanthomonas* pathogens like Xtt was MLSA (Young et al. 2008; Curland et al. 2018). Our ANI analyses, however, are not in agreement with the previous MLSA groupings (Fig. S1). Briefly, for MLSA, the sequences of four housekeeping genes *rpoD, dnaK, fyuA* and *gyrB* were concatenated according to Curland et al. (2018). The webtool NGPhylogeny was used to align the concatenated sequences with MAFFT, curate them with Gblocks and infer a tree with MrBayes (Lemoine et al. 2019). A tree with the studied Xtt strains and those from Curland et al. (2018) was created (Fig. S1) along with a tree only including studied strains for comparison to ANI-based analyses (Fig. S2). In agreement with previous work, the MLSA trees divided Xtt into 3 groups, one of which also contained the Xtu strain LW16 (Fig. S1, S2). Previous genomic studies had shown that Xtu was phylogenetically distinct based on whole genome analysis (Peng et al. 2016) or MLSA with 12 housekeeping genes (Langlois et al. 2017), suggesting that MLSA did not appropriately capture genetic diversity. Overall, our whole genome sequencing approach agrees with this finding because we find a phylogenetic separation between Xtt and Xtu and phylogenetic relationships between Xtt strains that do not match MLSA analysis. Therefore, four gene MLSA is not an appropriate method to define genetic classifications for Xtt.

*Xanthomonas* phytopathogens deploy a wide range of virulence factors, including secreted effectors, during pathogenesis (Timilsina et al. 2020). These effectors support pathogen nutrient acquisition and evasion of host defenses, but their recognition by a host plant can also trigger resistance to xanthomonads (Schornack et al. 2006; Lolle et al. 2020; Thomas et al. 2020).Carbohydrate active enzymes (CAZymes) are vital for plant-associated microbes to gain energy from the plant environment in which carbohydrate photosynthetic products are the main carbon source (Zhang et al. 2018). Some of these CAZymes are secreted and their identification can provide information about how a bacterium behaves and gains energy from its host. For example, one Type II secreted CAZyme, CbsA, functions as a key genetic determinant for tissue-specific adaptation between Xtt and Xtu (Gluck-Thaler et al. 2020). Because of the link between the presence of this gene and a pathovar-specific phenotype, we are now developing a subgroup-specific diagnostic.

Type III-secreted effectors (T3SEs) are directly injected into and manipulate host cells, often contributing to virulence (Rossier et al. 1999). One type of T3SE are transcription activator like effectors (TALEs). TALEs directly interact with specific host DNA sequences, with repetitive amino acid sequences that differ only in pairs of amino acids called repeat variable diresidues (RVDs) and promote transcription of downstream genes (Boch and Bonas 2010). This host manipulation frequently makes them major virulence factors in *Xanthomonas* pathogenesis (Perez-Quintero and Szurek 2019). For example, Xtu TALEs have a significant role in virulence on wheat. Little is known about the function of Xtt TALEs, although Xtt strains have approximately 5-8 TALEs according to southern blotting analysis of Iranian *X. translucens* (Khojasteh et al. 2020) and published genome sequences (Roman-Reyna et al. 2020; Shah et al. 2021). Our understanding of TALEs and their composition is limited despite increasing availability of *Xanthomonas* genomes, because long read sequencing is necessary to correctly describe and map highly repetitive TALEs in *Xanthomonas* genomes (Peng et al. 2016).

We predicted secreted proteins with SignalP 5.0 (Almagro Armenteros et al. 2019). Putative CAZymes were identified with dbCAN2 using the HMMER, DIAMOND and Hotpep algorithms (Zhang et al. 2018). T3SEs were identified using the BLAST 2.8.1+ blastx algorithm (Zhang et al. 2000) with studied genomes as queries and a database of known *Xanthomonas* T3SEs (xanthomonas.org), excluding TALEs which were analyzed separately (below). The BLAST results were filtered to include only hits with a coverage of over 200 amino acids and a percent amino acid identity of 60% or greater. TALEs in the eight studied genomes were identified and classified with AnnoTALE version 1.5 (Grau et al. 2016). FuncTAL version 1.1 in the QueTAL suite (Pérez-Quintero et al. 2015) was used to analyze the differences in TALE RVD patterns.

Though Xtt strains show a high degree of homology at the whole genome level, we hypothesized that their virulence factor complements would be variable in response to varying evolutionary pressures. To test this hypothesis, we identified and compared virulence factors in long-read genomes. All eight genomes each encoded more than 700 proteins with a predicted signal peptide (Table 1). Putative CAZymes were numerous and diverse, ranging in number from 112-117 per strain and representing a mix of glucoside hydrolases, glycosyltransferases, polysaccharide lyases and carbohydrate esterases (Table S2). This diversity likely underpins the ability of Xtt strains to exploit the complex carbohydrate environment of a host barley plant.

There are examples of highly conserved T3SEs. All studied strains possess copies of 15 putative T3SEs, suggesting that these proteins have conserved roles in Xtt pathogenesis (Fig. 2; Table S3). TALE distribution is more complex in comparison to the high conservation of other T3SEs in Xtt. Several TALEs are highly conserved according to their RVD patterns. For example, every tested strain possesses a TalCT and TalCV effector with identical RVD patterns (Fig. 3; Table S4). Eight TALEs were present in at least one Iranian and U.S. strain (Fig. 2). Despite the geographic separation and use of different barley varieties (Izadi et al. 2014; Mortazavian et al. 2014; Zhou et al. 2020), the identical RVD patterns in some TALEs suggests conserved roles in host manipulation (Fig. 3). Such conserved effectors, if they are critical for pathogenesis, are ideal elicitors to discover for broad-spectrum resistance in barley.

**Figure 2.**
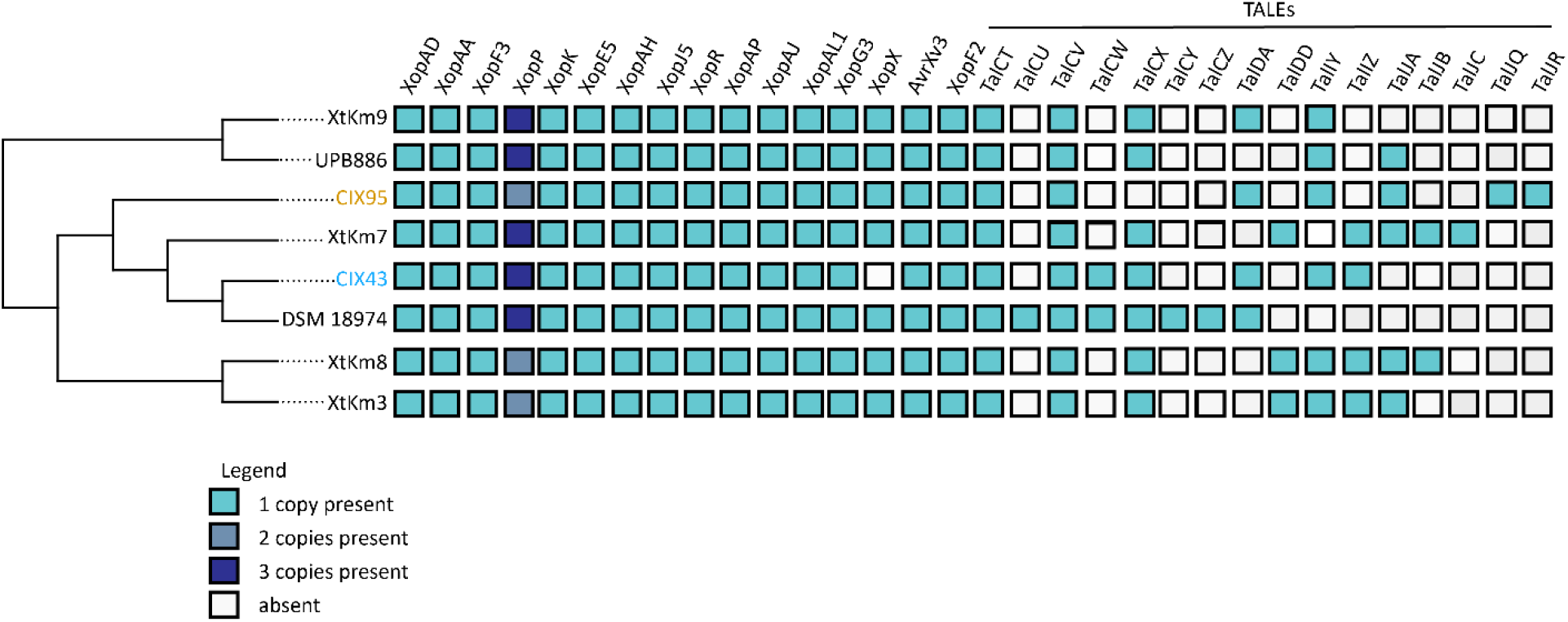
Xtt TALEs are diverse but other T3SEs are conserved. Putative T3SEs for eight long-read *X. translucens pv. translucens* genome assemblies were identified with a local Blastx against a database of known *Xanthomonas* effectors. Blue colored boxes represent the presence of a putative effector with shading representing the number of copies present. TALEs were identified and classified according to AnnoTALE (Grau et al. 2016) and their names begin with “Tal”. Whole genome ANI was calculated displayed strains and a tree was generated with the webtool Enveomics using the UPGMA clustering method (Rodriguez-R and Konstantinidis 2016).

**Figure 3.**
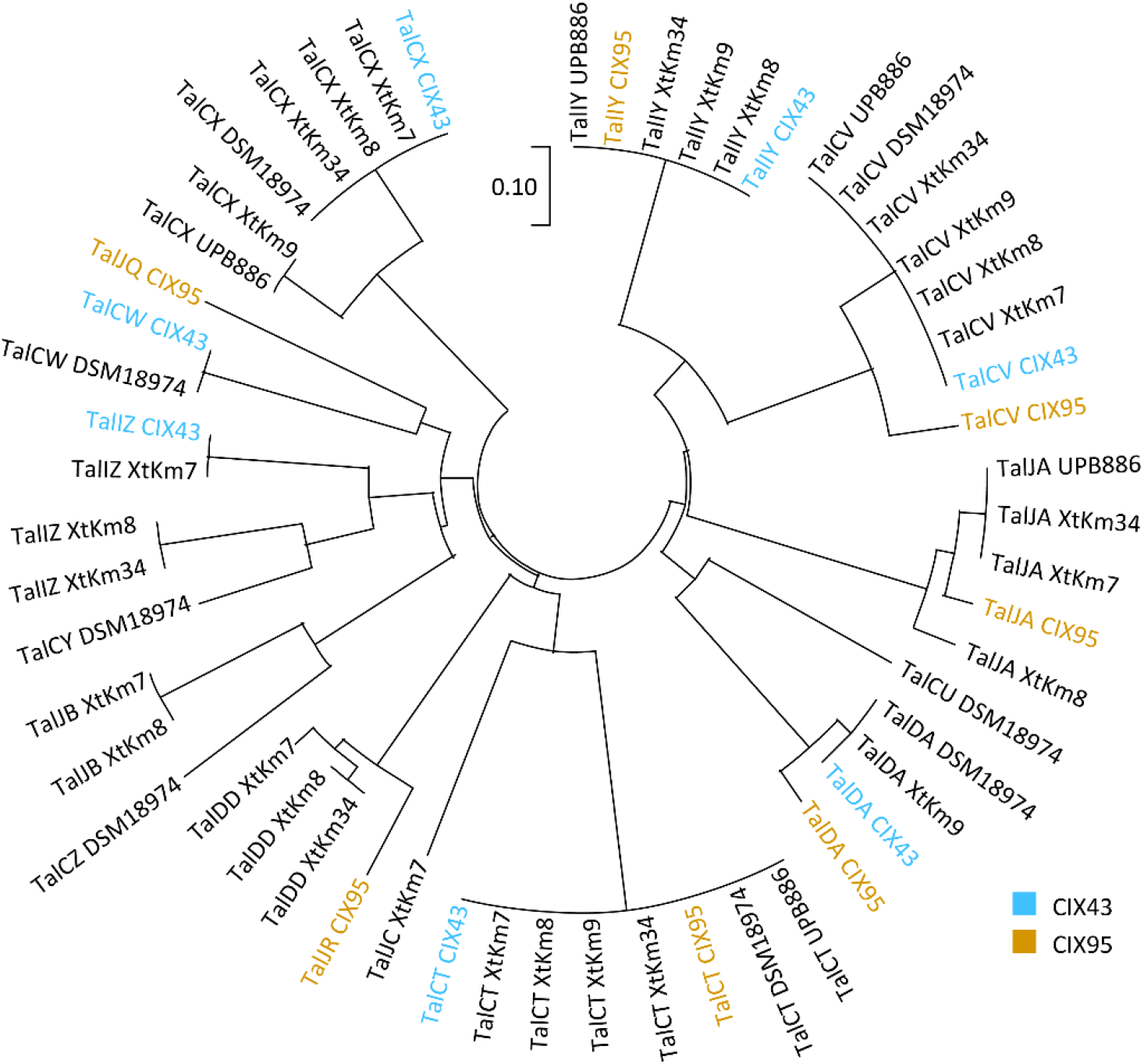
CIX95 has multiple unique TALEs. The colored names represent TALEs from the strains CIX43 (blue) and CIX95 (gold), sequenced in this study. The output was created with FuncTAL from the QueTAL suite of tools (Pérez-Quintero et al. 2015), using RVDs determined by AnnoTALE (Grau et al. 2016).

In contrast, there is large variability in the repertoire of TALEs that a particular strain possesses. According to their RVD patterns, 10 TALE classes identified in our analysis are present in multiple strains while six are unique to a single strain (Fig. 2; Fig. 3). These included two distinct TALEs in CIX95 in the classes TalJQ and TalJR. DSM 18974, XtKm7 and XtKm8 also include at least one TALE that does not match any others in the tested strains. The high diversity of TALE repertoires presents a challenge for breeders who attempt to characterize barley resistance against a limited panel of Xtt strains that may not represent the virulence capabilities of a field population.

In conclusion, we determined that globally, *X. translucens* pv. translucens strains, including CIX43 and CIX95, are highly genetically similar with three groups present in both Asia and North America. The *X. translucens* pv. translucens strains CIX43 and CIX95 are within the same subgroup and therefore more closely related than was previously suggested by MLSA. There are virulence factors that are highly conserved at the local and global levels, such as the TalCT and TalCV TALEs and 15 other T3SEs. Although their importance in virulence remains to be investigated, these effectors are potential targets for durable and broad resistance. On the other hand, there are diverse virulence factors at the population level, especially TALEs, such that resistance to one strain of *X. translucens* pv. translucens likely does not guarantee resistance to others. Based on the distinct virulence factor profiles observed in our small panel, multiple distinct strains should be included when completing host resistance phenotyping to increase the chances that discoveries are relevant to the field population.

Genetic resistance to bacterial leaf streak is lacking in elite malting barley varieties and has not been characterized for reaction to X. translucens. To develop representative phenotyping tests to screen germplasm, it is important to understand the diversity of the causal agent. The genomes of *X. translucens* pv. translucens strains CIX43 and CIX95 advance our knowledge about the *X. translucens* pv. translucens population in the Americas and are a resource relevant to control measures and barley breeding for cultivation. Representatives from all the Xtt LINgroups are already globally dispersed. Therefore, international regulation of seed is unlikely crucial for the control of pathogen spread.

## Acknowledgements

We are thankful to Drs. Adam Bogdanove, Stephen P. Cohen, and Ralf Koebnik for critical reading of earlier versions of this manuscript. We thank Drs. Jan Grau and Alvaro Pérez-Quintero for assistance in classifying TALEs and their RVD patterns. The authors acknowledge support from USDA National Institute of Food and Agriculture Award Number: 2018-67013-28490 through the NSF/NIFA Plant Biotic Interactions Program; the Ohio Department of Agriculture Specialty Crops Block Grant Number AGR-SCG-19-03; The American Malting Barley Association to JMJ; and an Environmental Fellowship from The Ohio State University College of Food, Agriculture and Environmental Science to NH.

## References

Almagro Armenteros, J. J., Tsirigos, K. D., Sønderby, C. K., Petersen, T. N., Winther, O., Brunak, S., et al. 2019. SignalP 5.0 improves signal peptide predictions using deep neural networks. Nat. Biotechnol. 37:420–423.

Boch, J., and Bonas, U. 2010. Xanthomonas AvrBs3 family-type III effectors: Discovery and function. Annu. Rev. Phytopathol. 48:419–436.

Bragard, C., Singer, E., Alizadeh, A., Vauterin, L., Maraite, H., and Swings, J. 1997. Xanthomonas translucens from small grains: Diversity and phytopathological relevance. Phytopathology. 87:1111–1117.

Curland, R. D., Gao, L., Bull, C. T., Vinatzer, B. A., Dill-Macky, R., Van Eck, L., et al. 2018. Genetic diversity and virulence of wheat and barley strains of Xanthomonas translucens from the upper midwestern United States. Phytopathology. 108:443–453.

Curland, R. D., Gao, L., Hirsch, C. D., and Ishimaru, C. A. 2020. Localized Genetic and Phenotypic Diversity of Xanthomonas translucens Associated With Bacterial Leaf Streak on Wheat and Barley in Minnesota. Phytopathology. 110:257–266.

Forster, R. L., and Schaad, N. W. 1988. Control of Black Chaff of Wheat with Seed Treatment and a Foundation Seed Health Program. Plant Dis. 72:935–938.

Gluck-Thaler, E., Cerutti, A., Perez-Quintero, A. L., Butchacas, J., Roman-Reyna, V., Madhavan, V. N., et al. 2020. Repeated gain and loss of a single gene modulates the evolution of vascular plant pathogen lifestyles. Sci. Adv. 6:4516–4529.

Goettelmann, F., Roman-Reyna, V., Cunnac, S., Jacobs, J. M., Bragard, C., Studer, B., et al. 2022. Complete genome assemblies of all Xanthomonas translucens pathotype strains reveal three genetically distinct clades. Front. Microbiol. 12:4386.

Grau, J., Reschke, M., Erkes, A., Streubel, J., Morgan, R. D., Wilson, G. G., et al. 2016. AnnoTALE: bioinformatics tools for identification, annotation and nomenclature of TALEs from Xanthomonas genomic sequences. Sci. Rep. 6:21077.

Habibian, M., Alizadeh Aliabadi, A., Hayati, J., and Rahimian, H. 2021. Investigation of the phenotypic and genetic diversity of Xanthomonas translucens pathovars, the causal agents of bacterial leaf streak of wheat and barley in parts of Iran. Plant Prot. (Scientific J. Agric.) 44:33–50.

Izadi, M. H., Rabbani, J., Emam, Y., Pessarakli, M., and Tahmasebi, A. 2014. Effects of salinity stress on physiological performance of various wheat and barley cultivars. J. Plant Nutr. 37:520–531.

Jaenicke, S., Bunk, B., Wibberg, D., Spröer, C., Hersemann, L., Blom, J., et al. 2016. Complete genome sequence of the barley pathogen Xanthomonas translucens pv. translucens DSM 18974T (ATCC 19319T). Genome Announc. 4:e01334–16.

Jones, L. R., Johnson, A. G., and Reddy, C. S. 1917. Bacterial-blight of barley. J. Agric. Res. 11:625–643.

Khojasteh, M., Shah, S. M. A., Haq, F., Xu, X., Taghavi, S. M., Osdaghi, E., et al. 2020. Transcription Activator-Like Effectors Diversity in Iranian Strains of Xanthomonas translucens. Phytopathology. 110:758–767.

Kolmogorov, M., Yuan, J., Lin, Y., and Pevzner, P. A. 2019. Assembly of long, error-prone reads using repeat graphs. Nat. Biotechnol. 37:540–546.

Langlois, P. A., Snelling, J., Hamilton, J. P., Bragard, C., Koebnik, R., Verdier, V., et al. 2017. Characterization of the Xanthomonas translucens Complex Using Draft Genomes, Comparative Genomics, Phylogenetic Analysis, and Diagnostic LAMP Assays. Phytopathology. 107:519–527.

Lemoine, F., Correia, D., Lefort, V., Doppelt-Azeroual, O., Mareuil, F., Cohen-Boulakia, S., et al. 2019. NGPhylogeny.fr: new generation phylogenetic services for non-specialists. Nucleic Acids Res. 47:W260–W265.

Lolle, S., Stevens, D., and Coaker, G. 2020. Plant NLR-triggered immunity: from receptor activation to downstream signaling. Curr. Opin. Immunol. 62:99–105.

Mortazavian, S. M. M., Nikkhah, H. R., Hassani, F. A., Sharif-Al-Hosseini, M., Taheri, M., and Mahlooji, M. 2014. GGE Biplot and AMMI Analysis of Yield Performance of Barley Genotypes across Different Environments in Iran. J. Agric. Sci. Technol. 16:609–622.

Overbeek, R., Olson, R., Pusch, G. D., Olsen, G. J., Davis, J. J., Disz, T., et al. 2014. The SEED and the Rapid Annotation of microbial genomes using Subsystems Technology (RAST). Nucleic Acids Res. 42:D206–D214.

Peng, Z., Hu, Y., Xie, J., Potnis, N., Akhunova, A., Jones, J., et al. 2016. Long read and single molecule DNA sequencing simplifies genome assembly and TAL effector gene analysis of Xanthomonas translucens. BMC Genomics. 17:21.

Pérez-Quintero, A. L., Lamy, L., Gordon, J. L., Escalon, A., Cunnac, S., Szurek, B., et al. 2015. QueTAL: a suite of tools to classify and compare TAL effectors functionally and phylogenetically. Front. Plant Sci. 6:545.

Perez-Quintero, A. L., and Szurek, B. 2019. A Decade Decoded: Spies and Hackers in the History of TAL Effectors Research. Annu. Rev. Phytopathol. 57:459–481.

Pesce, C., Bolot, S., Cunnac, S., Portier, P., Saux, M. F. Le, Jacques, M. A., et al. 2015. High-quality draft genome sequence of the Xanthomonas translucens pv. cerealis pathotype strain CFBP 2541. Genome Announc. 3:e01574–14.

Rodriguez-R, L. M., and Konstantinidis, K. T. 2016. The enveomics collection: a toolbox for specialized analyses of microbial genomes and metagenomes. PeerJ Prepr. 4:e1900v1.

Roman-Reyna, V., Luna, E. K., Pesce, C., Vancheva, T., Chang, C., Ziegle, J., et al. 2020. Genome resource of barley bacterial blight and leaf streak pathogen Xanthomonas translucens pv. translucens strain UPB886. Plant Dis. 104:13–15.

Rossier, O., Wengelnik, K., Hahn, K., and Bonas, U. 1999. The Xanthomonas Hrp type III system secretes proteins from plant and mammalian bacterial pathogens. Proc. Natl. Acad. Sci. U. S. A. 96:9368–9373.

Sapkota, S., Mergoum, M., and Liu, Z. 2020. The translucens group of Xanthomonas translucens: Complicated and important pathogens causing bacterial leaf streak on cereals. Mol. Plant Pathol. 21:291–302.

Schornack, S., Meyer, A., Römer, P., Jordan, T., and Lahaye, T. 2006. Gene-for-gene-mediated recognition of nuclear-targeted AvrBs3-like bacterial effector proteins. J. Plant Physiol. 163:256–272.

Shah, S. M. A., Khojasteh, M., Wang, Q., Taghavi, S. M., Xu, Z., Khodaygan, P., et al. 2021. Genomics-Enabled Novel Insight Into the Pathovar-Specific Population Structure of the Bacterial Leaf Streak Pathogen Xanthomonas translucens in Small Grain Cereals. Front. Microbiol. 12:1265.

Tambong, J. T., Xu, R., Gerdis, S., Daniels, G. C., Chabot, D., Hubbard, K., et al. 2021. Molecular Analysis of Bacterial Isolates From Necrotic Wheat Leaf Lesions Caused by Xanthomonas translucens, and Description of Three Putative Novel Species, Sphingomonas albertensis sp. nov., Pseudomonas triticumensis sp. nov. and Pseudomonas foliumensis sp. nov. Front. Microbiol. 12.

Tatusova, T., Dicuccio, M., Badretdin, A., Chetvernin, V., Nawrocki, E. P., Zaslavsky, L., et al. 2016. NCBI prokaryotic genome annotation pipeline. Nucleic Acids Res. 44:6614–6624.

Thomas, N. C., Hendrich, C. G., Gill, U. S., Allen, C., Hutton, S. F., and Schultink, A. 2020. The Immune Receptor Roq1 Confers Resistance to the Bacterial Pathogens Xanthomonas, Pseudomonas syringae, and Ralstonia in Tomato. Front. Plant Sci. 11:463.

Tian, L., Huang, C., Mazloom, R., Heath, L. S., and Vinatzer, B. A. 2020. LINbase: a web server for genome-based identification of prokaryotes as members of crowdsourced taxa. Nucleic Acids Res. 48:W529–W537.

Timilsina, S., Potnis, N., Newberry, E. A., Liyanapathiranage, P., Iruegas-Bocardo, F., White, F. F., et al. 2020. Xanthomonas diversity, virulence and plant–pathogen interactions. Nat. Rev. Microbiol. 18:415–427.

Young, J. M., Park, D.-C., Shearman, H. M., and Fargier, E. 2008. A multilocus sequence analysis of the genus Xanthomonas. Syst. Appl. Microbiol. 31:366–377.

Zhang, H., Yohe, T., Huang, L., Entwistle, S., Wu, P., Yang, Z., et al. 2018. dbCAN2: A meta server for automated carbohydrate-active enzyme annotation. Nucleic Acids Res. 46:W95–W101.

Zhang, Z., Schwartz, S., Wagner, L., and Miller, W. 2000. A Greedy Algorithm for Aligning DNA Sequences. J. Comput. Biol. 7:203–214.

Zhou, B., Jin, Z., Schwarz, P., and Li, Y. 2020. Impact of Genotype, Environment, and Malting Conditions on the Antioxidant Activity and Phenolic Content in US Malting Barley. Fermentation. 6:48.

