## Supplemental Materials for "Comparative genomics of Minnesotan barley-infecting *Xanthomonas translucens* shows overall genomic similarity but virulence factor diversity"

Supplemental Figures

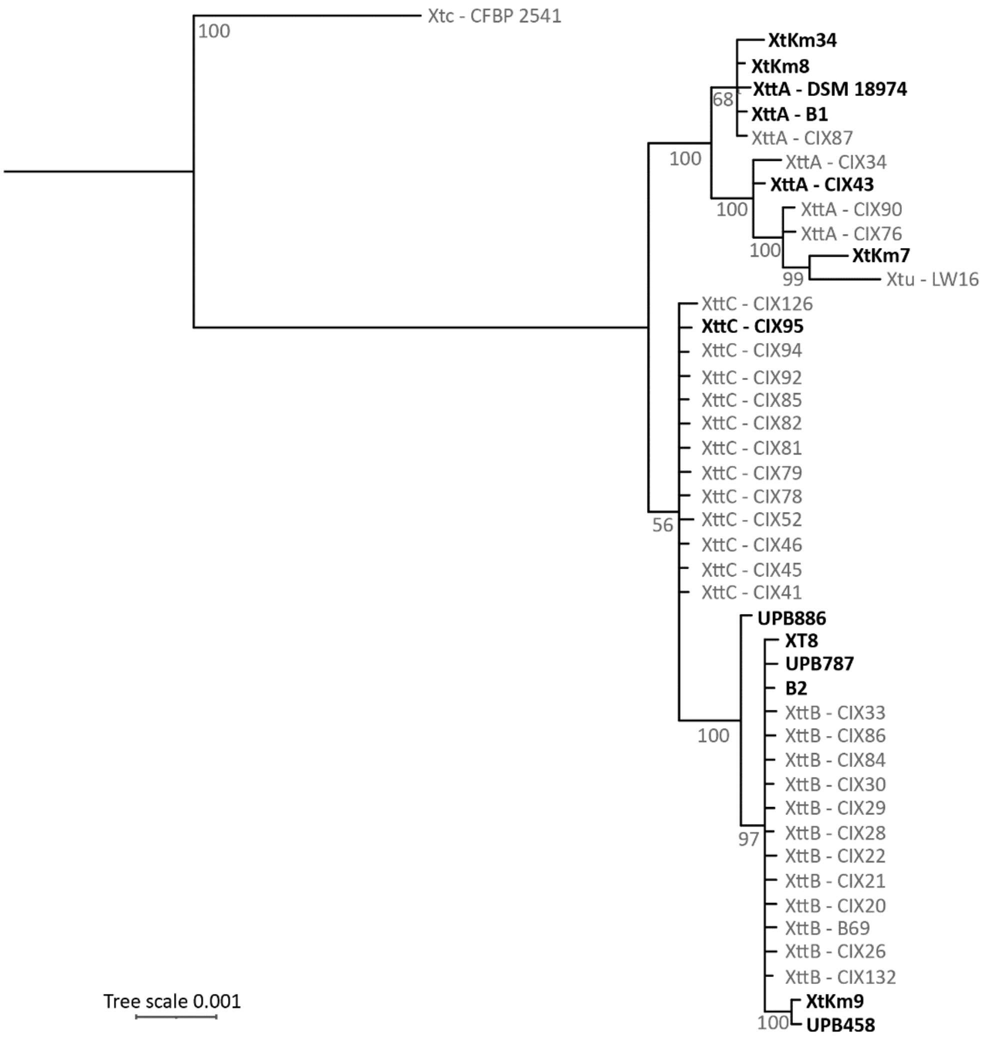


Supplemental Figure 1. Xtt strains separate into three groups according to MLSA and Xtt group A strains cluster with Xtu strain LW16. Housekeeping genes *dyaK*, *fyuA*, *gyrB* and *rpoD* were concatenated for Xtt strains from Curland et al. 2018 and Xtt strains included in this study (bold). Strains previously placed into groups are labeled as XttA, XttB or XttC. Using the webtool NGPhylogeny, the concatenated sequences were aligned with MAFFT, curated with Gblocks and a tree was inferred with MrBayes (Lemoine et al. 2019). Posterior probability percentages of branches are displayed. The tree was rooted on CFBP 2541.

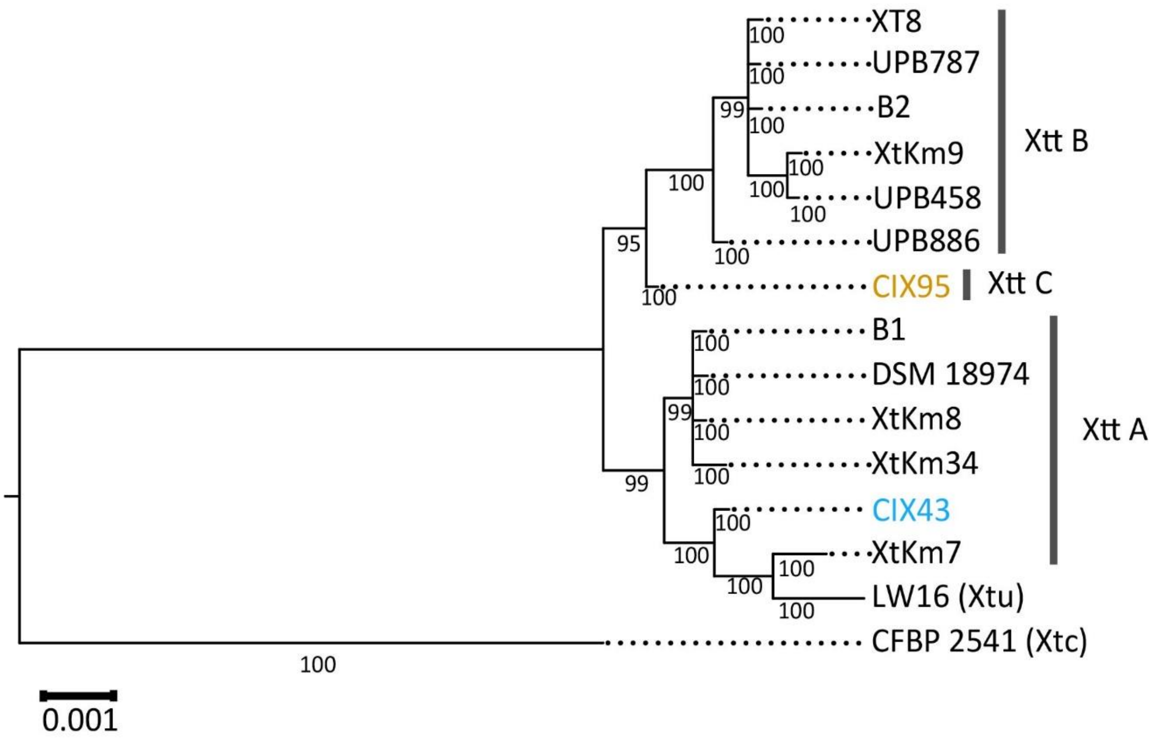


Supplemental Figure 2. MLSA does not match with whole genome ANI analysis. Housekeeping genes *dyaK*, *fyuA*, *gyrB* and *rpoD* were concatenated for all publicly available *X. translucens* pv. translucens strains and the outgroup strains *X. translucens* pv. undulosa LW16 and *X. translucens* pv. cerealis CFBP 2541. Using the webtool NGPhylogeny, the concatenated sequences were aligned with MAFFT, curated with Gblocks and a tree was inferred with MrBayes (Lemoine et al. 2019). Posterior probability percentages of branches are displayed. The tree was rooted on Xtc CFBP 2541.
